# Thioproline formation as a driver of formaldehyde toxicity in *Escherichia coli*

**DOI:** 10.1101/2020.03.19.981027

**Authors:** Jenelle A. Patterson, Hai He, Jacob S. Folz, Qiang Li, Mark A. Wilson, Oliver Fiehn, Steven D. Bruner, Arren Bar-Even, Andrew D. Hanson

## Abstract

Formaldehyde (HCHO) is a reactive carbonyl compound that formylates and cross-links proteins, DNA, and small molecules. It is of specific concern as a toxic intermediate in the design of engineered pathways involving methanol oxidation or formate reduction. The high interest in engineering these pathways is not, however, matched by engineering-relevant information on precisely why HCHO is toxic or on what damage-control mechanisms cells deploy to manage HCHO toxicity. The only well-defined mechanism for managing HCHO toxicity is formaldehyde dehydrogenase-mediated oxidation to formate, which is counterproductive if HCHO is a desired pathway intermediate. We therefore sought alternative HCHO damage-control mechanisms via comparative genomic analysis. This analysis associated homologs of the *Escherichia coli pepP* gene with HCHO-related one-carbon metabolism. Furthermore, deleting *pepP* increased the sensitivity of *E. coli* to supplied HCHO but not other carbonyl compounds. PepP is a proline aminopeptidase that cleaves peptides of the general formula X-Pro-Y, yielding X + Pro-Y. HCHO is known to react spontaneously with cysteine to form the close proline analog thioproline (thiazolidine-4-carboxylate), which is incorporated into proteins and hence into proteolytic peptides. We therefore hypothesized that thioproline-containing peptides are toxic and that PepP cleaves these aberrant peptides. Supporting this hypothesis, PepP cleaved the model peptide Ala-thioproline-Ala as efficiently as Ala-Pro-Ala *in vitro* and *in vivo*, and deleting *pepP* increased sensitivity to supplied thioproline. Our data thus (i) provide biochemical genetic evidence that thioproline formation contributes substantially to HCHO toxicity and (ii) make PepP a candidate damage-control enzyme for engineered pathways having HCHO as an intermediate.

## Introduction

Formaldehyde (HCHO) is a highly reactive, toxic, and mutagenic carbonyl compound that all organisms produce in minor amounts during normal metabolism [1-3]. It is also a major and mandatory intermediate in natural and engineered (‘synthetic’) methylotrophic and formatotrophic pathways [1,4-6], and in synthetic pathways to which methanol supplies reducing power but not carbon [7] (Figure 1).

**Figure 1.**
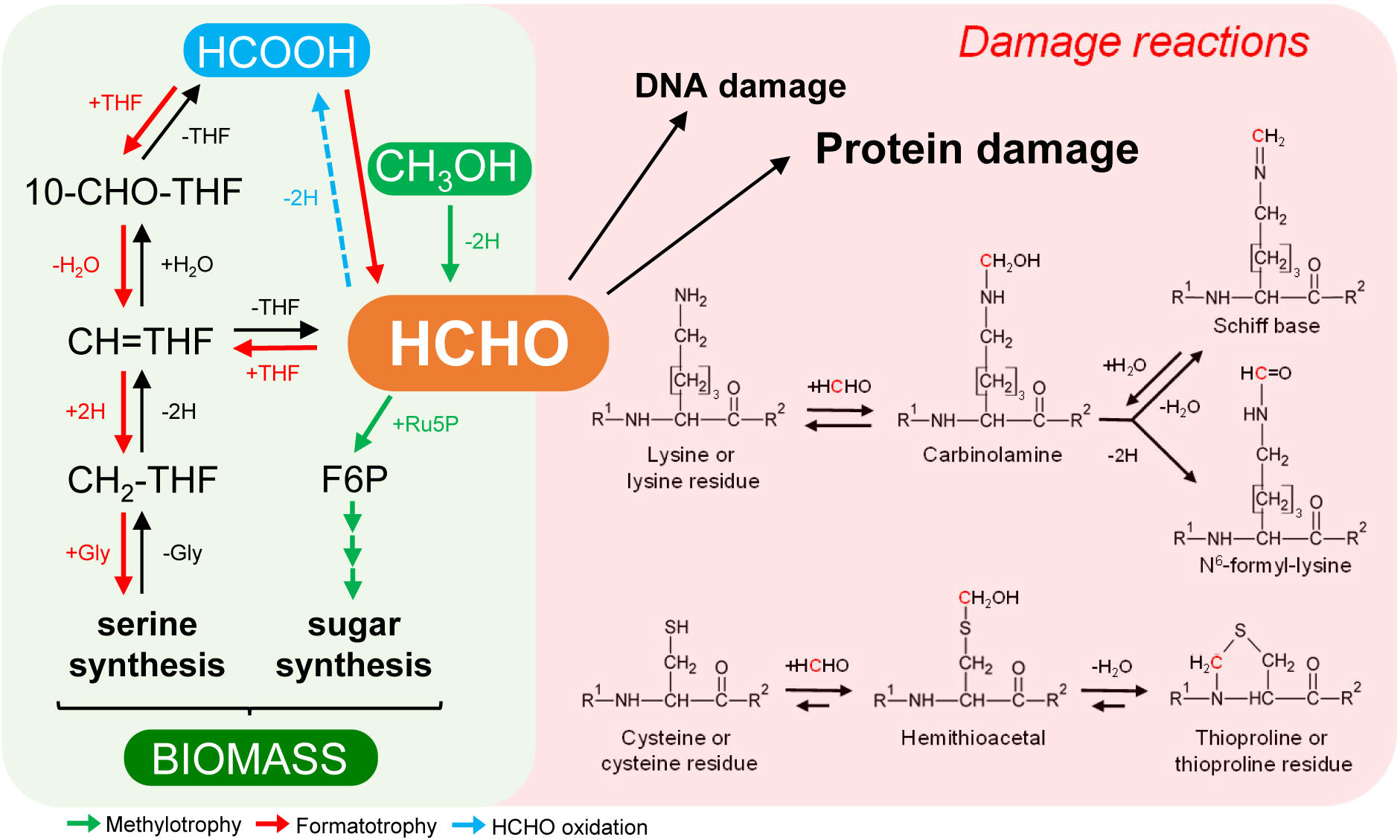
Formaldehyde as a central one-carbon metabolite and driver of damage reactions. Formaldehyde (HCHO) participates in methylotrophy and formatotrophy pathways but reacts spontaneously with amino and thiol groups, e.g. those in the side chains of lysine and cysteine, respectively. The Schiff base formed with lysine undergoes various cross-linking reactions with proteins and nucleic acids. HCHO can be detoxified by oxidation to formate (HCOOH) but this route is incompatible with methylotrophy or formatotrophy pathways in which HCHO is an obligatory intermediate. R^1^, H or peptide chain; R^2^, OH or peptide chain; Ru5P, ribulose 5-phosphate; THF, tetrahydrofolate, 10-CHO-, 10-formyl; CH=, 5,10-methenyl; CH_2_-, 5,10-methylene.

The toxicity and mutagenicity of HCHO lie in its ability to react spontaneously with amino and thiol groups. With amines such as the 6-amino group of peptidyl lysine, HCHO forms a carbinolamine that can either be oxidized to an *N*-formyl derivative or undergo dehydration, yielding a labile Schiff base that can form cross-links (methylene bridges) with various amino acids and nucleobases [8,9] (Figure 1). With thiols, notably free and peptidyl cysteine, HCHO forms the cyclic derivative thioproline (thiazolidine-4-carboxylate) [10,11] (Figure 1). All of these reactions except oxidation and cross-linking are reversible, with the HCHO condensation direction being thermodynamically preferred. Although it is clear that the above general types of damage reactions underlie the toxic and mutagenic effects of HCHO, the specific reactions that actually drive these effects and their relative importance *in vivo* remain poorly defined [10]. The least poorly defined toxic effect is probably the *N*^6^-formylation of lysine residues in certain histones, which is inferred to impair histone function [12].

The archetypal defense against HCHO toxicity is to oxidize HCHO to non-toxic formate via glutathione-dependent or -independent dehydrogenases [1-3] (Figure 1). This detoxification strategy is counter-productive if HCHO is a pathway substrate, as in methylotrophic or formatotrophic pathways, because it leads to loss of the substrate. Cells operating methylotrophic or formatotrophic pathways must therefore cope with HCHO because they cannot simply detoxify it by dehydrogenation [13]. A major coping mechanism is fine-tuning the expression of HCHO-producing and -consuming enzymes [13,14], but this delicate flux-balancing act entails non-negligible steady-state HCHO levels as well as possible HCHO spikes due to upshifts in methanol availability. It is consequently worth identifying other mechanisms that cells use to control HCHO damage. The only known damage-control mechanism of this type appears to be deformylation of *N*^6^-formyl lysine residues by bacterial and mammalian sirtuins [15].

The research we report here was based on two ideas: that there are HCHO damage-control enzymes left to discover [16] and that such enzymes are valuable components to have in hand when designing and building metabolic pathways that involve HCHO [17]. After applying comparative genomics to identify a candidate HCHO damage-control enzyme, we obtained genetic and biochemical evidence that *Escherichia coli* PepP, an Xaa-Pro aminopeptidase, mitigates downstream consequences of the presence of HCHO-generated thioproline residues in proteins. That deleting *pepP* increases HCHO sensitivity provides genetic evidence that thioproline formation is one of the drivers of HCHO toxicity.

## Materials and methods

### Bioinformatics

Comparative genomics analyses used STRING [18] (https://string-db.org/) and SEED [19] (http://pubseed.theseed.org/) databases and tools. The guide genes used to search for candidate HCHO damage-control genes were those mediating reactions near HCHO in KEGG maps 00680 (methane metabolism) and 00670 (one carbon pool by folate). Full results of the SEED analysis are encoded in the SEED subsystem named ‘PepP-C1’. Phylogenetic trees were drawn with MEGA6 [20].

### Chemicals

Phusion® High-Fidelity DNA polymerase (New England Biolabs, Ipswich, MA) was used for PCR reactions. All other chemicals were from Sigma–Aldrich, unless otherwise stated. HCHO solution (37%, with methanol stabilizer) was from Fisher Scientific (Cat. No. BP531-500). Tripeptides used for *in vitro* assays and growth experiments (H-Ala-Pro-Ala-OH, H-Ala-Thz-Ala-OH trifluoroacetate, and H-Glu(Thz-Gly-OH)-OH trifluoroacetate) were synthesized by Bachem (UK) Ltd. (St. Helens, UK); their purities (>98%) were validated by HPLC. *N*^6^-Formyl-L-lysine was from TCI Chemicals (Cat. No. F0136). Formylated Boc-L-Lys-AMC was synthesized from Boc-L-Lys-AMC (Peptide Solutions LLC, Tucson, AZ, Cat. No. BAA3650.0100) using a modification of a published procedure [21]. To a solution of Boc-L-Lys-AMC (50 mg, 0.12 mmol, 1.0 equiv.) and a catalytic amount of DIPEA (0.01 mmol) in dry dimethylformamide (2 ml) was added 2,2,2-trifluoroethyl formate (61 mg, 0.48 mmol, 4.0 equiv.) and the reaction mixture was stirred at 22°C for 12 h. The mixture was concentrated *in vacuo* and purified by silica gel column chromatography (CH_2_Cl_2_:MeOH, 20/1 v/v) to give the product (36 mg, 67%). ^1^H NMR (300 MHz, DMSO-*d*_6_): *δ* 10.43 p.p.m. (s, 1H), 7.98 (s, 2H), 7.78 (d, 1H, *J* = 3 Hz), 7.73 (d, 1H, *J* = 9 Hz), 7.50 (d, 1H, *J* = 9 Hz), 7.14 (d, 1H, *J* = 9 Hz), 6.26 (s, 1H), 4.06 (m, 1H), 3.06 (m, 2H), 2.40 (s, 3H), 1.62-1.57 (m, 2H), 1.42-1.38 (m, 11H), 1.29-1.27 (m, 2H). LC-MS (ESI) m/z 430.2 [M - H] ^-^.

### Bacterial strains and expression constructs

Keio collection wild type background (BW25113), Δ*pepP*, and Δ*yhbO* strains [22] were the gift of V. de Crecy-Lagard (University of Florida). Primers are listed in Supplementary Table S1. All constructs were sequence-verified. For Δ*pepP* mutant complementation, the *E. coli* PepP coding sequence was PCR-amplified (primers 1 and 2) from BW25113 genomic DNA prepared using a GeneJET Genomic DNA Purification Kit (Thermo Scientific, Cat No. K0721) according to the manufacturer’s instructions. Amplicons were digested with *Eco*RI and *Sal*I (New England Biolabs) and ligated into pBAD24. P1 phage transduction of the Δ*yhbO*::Kan^r^ locus from the Keio Δ*yhbO* donor into the recipient BW25113 strain was performed according to [23]. P1 phage was isolated from strain EST877 P1cml, clr100 (the gift of L.N. Csonka, Purdue University), which was precultured in 2 ml Luria Broth (LB) with 30 μg/ml chloramphenicol at 30°C. The preculture was subcultured (1:100) in 50 ml LB and grown at 30°C with shaking until OD_600_ ∼0.8 before shifting the temperature to 40°C for 35 min, then 37°C for 1 h. The cells were lysed by adding 0.5 ml chloroform; after 10 min, the phage-containing supernatant was collected, filtered (0.45 μm) and stored at 4°C. Transduced cells were selected on M9 minimal medium plates containing 50 μg/ml kanamycin at 37°C for 48 h, and positive strains were sequence-verified.

### Media and culture conditions

Growth experiments were performed in M9 minimal medium (47.8 mM Na_2_HPO_4_, 22 mM KH_2_PO_4_, 8.6 mM NaCl, 18.7 mM NH_4_Cl, 2 mM MgSO_4_ and 100 μM CaCl_2_), supplemented with trace elements (134 μM EDTA, 31 μM FeCl_3_, 6.2 μM ZnCl_2_, 0.76 μM CuCl_2_, 0.42 μM CoCl_2_, 1.62 μM H_3_BO_3_, 0.081 μM MnCl_2_) and 22 mM glucose, unless otherwise specified. NH_4_Cl was omitted for the nitrogen source experiments. For liquid growth curves, strains were precultured in 4 ml M9 medium containing 10 mM glucose. The precultures were harvested and washed three times in M9 medium, then inoculated in M9 medium containing 10 mM glucose plus specified additions, with a starting OD_600_ of 0.005. One hundred and fifty μl of culture was added to each well of 96-well microplates (Nunclon Delta Surface, Thermo Scientific), then 50 μl mineral oil (Sigma-Aldrich) was added to each well to avoid evaporation (while enabling gas diffusion). The 96-well microplates were incubated at 37°C in a microplate reader (BioTek EPOCH 2). The shaking program cycle (controlled by Gen5 v3) had four shaking phases, lasting 60 s each: linear shaking followed by orbital shaking, both at an amplitude of 3 mm, then linear shaking followed by orbital shaking both at an amplitude of 2 mm. The OD_600_ in each well was monitored and recorded after every three shaking cycles (∼16.5 min). Raw data from the plate reader were calibrated to normal cuvette-measured OD_600_ values according to OD_cuvette_ = OD_plate_/0.23.

For HCHO insult experiments, 50 ml of M9 medium was inoculated with BW25113 and Δ*pepP* overnight precultures to a starting OD_600_ of 0.01 and incubated with shaking at 37°C until OD_600_ reached 0.2. Pilot experiments showed that adding HCHO to cultures to a final concentration of 1 mM caused a cessation of growth after which BW25113 and Δ*pepP* strains resumed growth after 13 h and 20 h, respectively. For untreated samples, cells were collected by centrifugation at OD_600_ of 0.2, flash frozen and stored at -80°C; HCHO-treated samples were similarly collected 2 h after the insult began.

M9 medium plates containing 1% agar were used to test growth on solid medium. When applicable, 400 μM formaldehyde (or 60 mM acetaldehyde, 5 mM glyoxal, or 0.5 mM methylglyoxal) were added to the medium after cooling to <45°C. Overnight M9 medium precultures were used to inoculate (1:100) 2-ml M9 medium cultures, which were grown at 37°C until OD_600_ of 1.0 before ten-fold serially diluting and spotting 3.5-μl aliquots on the plates. Plates were incubated at 37°C for 16-20 h before imaging.

To test Ala-Pro-Al or Ala-thioproline-Ala as sole nitrogen source, M9 plates were prepared as above except that NH_4_Cl was omitted. Overnight M9 medium precultures of BW25113 and Keio Δ*pepP* strains were used to inoculate (1:100) 10-ml M9 medium cultures, which were grown at 37°C until OD_600_ of ∼1.0. Cells from each strain were 10-fold diluted into water agar solution (0.75% w/v) cooled to ∼40°C, and 5 ml was overlaid on plates prewarmed to 37°C. Once cooled, 10-mm disks of sterile No. 2 Whatman filter paper were imbibed with 10 μl of 25 mg/ml alanine or 50 mg/ml tripeptide solution, then applied to the agar-overlaid plates. The plates were incubated at 37°C for 24 h before imaging.

### Production and purification of recombinant PepP

For PepP protein expression, the coding sequence was PCR-amplified (primers 3 and 4) from wild type BW25113 genomic DNA, adding an N-terminal His_6_-tag as well as 3’ and 5’ sequences complementary to the pET15b multiple cloning site. Following pET15b PCR amplification (primers 5 and 6), the vector and insert amplicons were gel-purified (GeneJet Gel Extraction Kit, Thermo Scientific Cat No. K0691) according to the manufacturer’s instructions and inserted using circular polymerase extension cloning [24]. To produce recombinant PepP protein, *E. coli* BL21 (DE3) RIPL harboring the expression plasmid was grown in 100 ml LB medium at 37°C until an OD_600_ of 0.8, after which the cultures were supplemented with 1.0 mM isopropyl-β-D-thiogalactoside and incubated a further 2 h at 22°C. Cells were collected by centrifugation, frozen in liquid N_2_, and stored at -80°C. Pellets were resuspended in 1 ml ice-cold lysis buffer (50 mM potassium phosphate, pH 8.0, 300 mM NaCl, and 10 mM imidazole), and sonicated (Fisher Scientific Ultrasonic Dismembrator, model 150E) using five 12-s pulses at 70% power, cooling on ice for 30 s between pulses. Following centrifugation (14 000 ***g***, 5 min, 4°C), the supernatant was applied to a 0.1 ml column of Ni^2+^-NTA Superflow resin (Qiagen Cat No. 30410) and protein was purified using the manufacturer’s protocol. The purified protein was desalted using a PD-25 column (GE Healthcare), eluting with 50 mM potassium phosphate, pH 7.5, containing 10% (v/v) glycerol; aliquots (50 μl) were frozen in liquid N_2_ and stored at -80°C. Protein concentration was determined by Bradford dye-binding assays [25].

### Enzyme assays

For all activity assays, PepP protein was pre-incubated with 0.5 mM MnCl_2_ for 15 min on ice. To assay deformylase activity, 40-μl reactions containing 2 μg PepP, 0.5 mM MnCl_2_, 50 mM potassium phosphate, pH 7.5, and 5 mM *N*^6^-formyl-L-lysine or 2 mM formylated BOC-Lys-AMC (diluted from a 30 mM stock solution in dimethylsulfoxide) were incubated at 22°C for 16 h. To quantify formate release, 0.05 units of formate dehydrogenase (Megazyme, Bray, Ireland, Cat. No. E-FDHCB) and 1 mM NAD were added (final reaction volume of 80 μl, 22°C). Absorbance at 340 nm for *N*^6^-formyl-L-lysine and 365 nm for formylated BOC-Lys-AMC was recorded every 15 s for 5 min. Dimethylsulfoxide at the concentration present in the assays had no effect on formate dehydrogenase.

Aminopeptidase activity against glutathione and γ-Glu-thioproline-Gly was assayed 22°C in 80-μl reactions containing 2 μg PepP, 0.5 mM MnCl_2_, 50 mM HEPES, pH 8.8, 1 mM NAD, and 0.02 units L-glutamate dehydrogenase (Sigma-Aldrich Cat. No. G2626). Reactions were started by adding 2 mM substrate. Assays using Ala-Pro-Ala and Ala-thiopro-Ala as substrate contained 2 μg PepP, 0.5 mM MnCl_2_, 50 mM Tris-HCl, pH 8.8, 1 mM NAD, and 0.05 units alanine dehydrogenase (Sigma-Aldrich Cat, No. 73063), and were started by adding 0.1-3 mM substrate. Absorbance at 340 nm was recorded every 10 s for 5 min for all aminopeptidase activity assays. Three independent assays at six substrate concentrations were performed to determine enzyme kinetics, which were calculated by fitting data to the Michaelis-Menten equation using Prism software (GraphPad Software, Inc., La Jolla, CA).

### LC-MS analysis of protein-bound N-formyl lysine

Cell pellets were resuspended in 1 ml PBS buffer (10 mM sodium phosphate, pH 7.4, 137 mM NaCl, 2.7 mM KCl), sonicated as above, and centrifuged to clear. Soluble proteins were precipitated in 80% (v/v) acetone and harvested by centrifugation (5000 ***g***, 3 min, 4°C); the pellets were air-dried overnight at 22°C and stored at -20°C. The pelleted proteins were mixed with 200 μl 1 M NaOH and heated at 50°C until dissolved (briefly vortexing as necessary) before adding 400 μl 100 mM ammonium bicarbonate buffer, pH 8.5 (brought to pH 8.5 with 5 M HCl). Samples were then desalted on PD-25 columns, eluting with 100 mM ammonium bicarbonate buffer, pH 8.5, and protein content was determined as above. To digest proteins to free amino acids, *Streptomyces griseus* protease [11] was added (5 μg per 50 μg protein) and the mixtures were incubated at 37°C for ∼16 h. The digests were vacuum-dried and stored at -20°C. Digests were dissolved by adding 0.5 ml HPLC grade 80:20 acetonitrile/water (v/v), vortexing for 30 s, sonicating for 5 min, and vortexing for another 30 s. From this solution 0.1 ml was aliquoted into a clean 1.5-ml microcentrifuge tube and dried under vacuum. Dried samples were resuspended in 0.04 ml of run solvent consisting of 80:20 acetonitrile/water (v/v) with deuterium labeled internal standards (IS) (Supplementary Table S2). To resuspend, solvent was added, samples were vortexed for 30 s, sonicated for 5 m, vortexed again for 30 s, centrifuged (16000 ***g***, 2 min, room temperature), and transferred to anLC-MS vial with glass insert. Samples were stored at 4°C in an autosampler and analyzed using a Vanquish LC system (Thermo Fisher Scientific), coupled to a 6500+ QTrap mass spectrometer (SCIEX) operated in multiple reaction monitoring (MRM) mode. Standard curves of analytes were diluted in LC-MS grade 80:20 acetonitrile/water (v/v), dried under vacuum, and resuspended in the same run solvent as samples. A Waters Acquity UPLC BEH Amide column (150 mm × 2.1 mm, 1.7 μm particle size) with an Acquity VanGuard BEH Amide pre-column (5 mm × 2.1 mm, 1.7 μm particle size) was held at 45°C throughout analysis. Mobile phase (A) was LC-MS grade water and (B) was 95:5 LC-MS grade acetonitrile/ water (v/v). Both mobile phases were modified to 10 mM ammonium formate and 0.125% formic acid. The mobile phase flow was 0.4 ml/min and the gradient was 100% B from 0-2 min, brought to 70% B between 2 and 7.7 min, brought to 40% B between 7.7 and 9.5 min, returned to 100% B between 9.5 and 12.75 min and maintained at 100% B until 17 min. Five µl of sample was injected and needle wash solution was 1:1 acetonitrile/water (v/v). Collision gas was set to “Medium”, IonSpray voltage was 5500 V and temperature was 250°C. Linear 5-7 point calibration curves of the ratio of analyte to IS with 1/x weighting were fit for lysine (IS: D8 lysine), *N*^6^-formyl lysine (IS: D5 threonine), and cystine (IS: D8 lysine) in MultiQuant version 3.0.3 (SCIEX). The background rate of *N*^6^-formylation of lysine during analysis was 0.03% determined from a calibration curve of pure lysine standard spiked into the matrix. Background formylation was corrected for by subtracting 0.0003*[lysine] from the measured concentrations of *N*^6^-formyl lysine in samples.

## Results and discussion

### Comparative genomics links PepP with one-carbon metabolism

We first used comparative genomics analysis [26] with the STRING [18] and SEED [19] databases to find candidate genes that could play a role in controlling HCHO damage. We searched for genes that are consistently clustered on the chromosome with genes of one-carbon metabolism, in which HCHO is central. These one-carbon metabolism ‘guide genes’ included genes encoding enzymes that produce or consume HCHO (e.g. methanol and formaldehyde dehydrogenases) and genes of tetrahydrofolate (THF)-mediated one-carbon metabolism, which is both a disposal route for HCHO and a potential HCHO source via spontaneous dissociation of 5,10-methylene-THF [27]. The analysis detected genes encoding homologs of *E. coli* aminopeptidase PepP clustered with genes encoding subunits of the glycine cleavage complex, which generates a THF-bound HCHO unit [28], and genes for 5-formyl-THF cyclo-ligase, which recycles a one-carbon folate byproduct of serine hydroxymethyltransferase, another enzyme that generates a THF-bound HCHO unit [29] (Figure 2A). These clustering arrangements are unlikely to be due to chance alone because they involve different HCHO-related genes and occur in bacteria from four different phyla. Moreover, the clustering specifically involves the PepP aminopeptidase family, but not related aminopeptidase families such as YpfF (Figure 2B).

**Figure 2.**
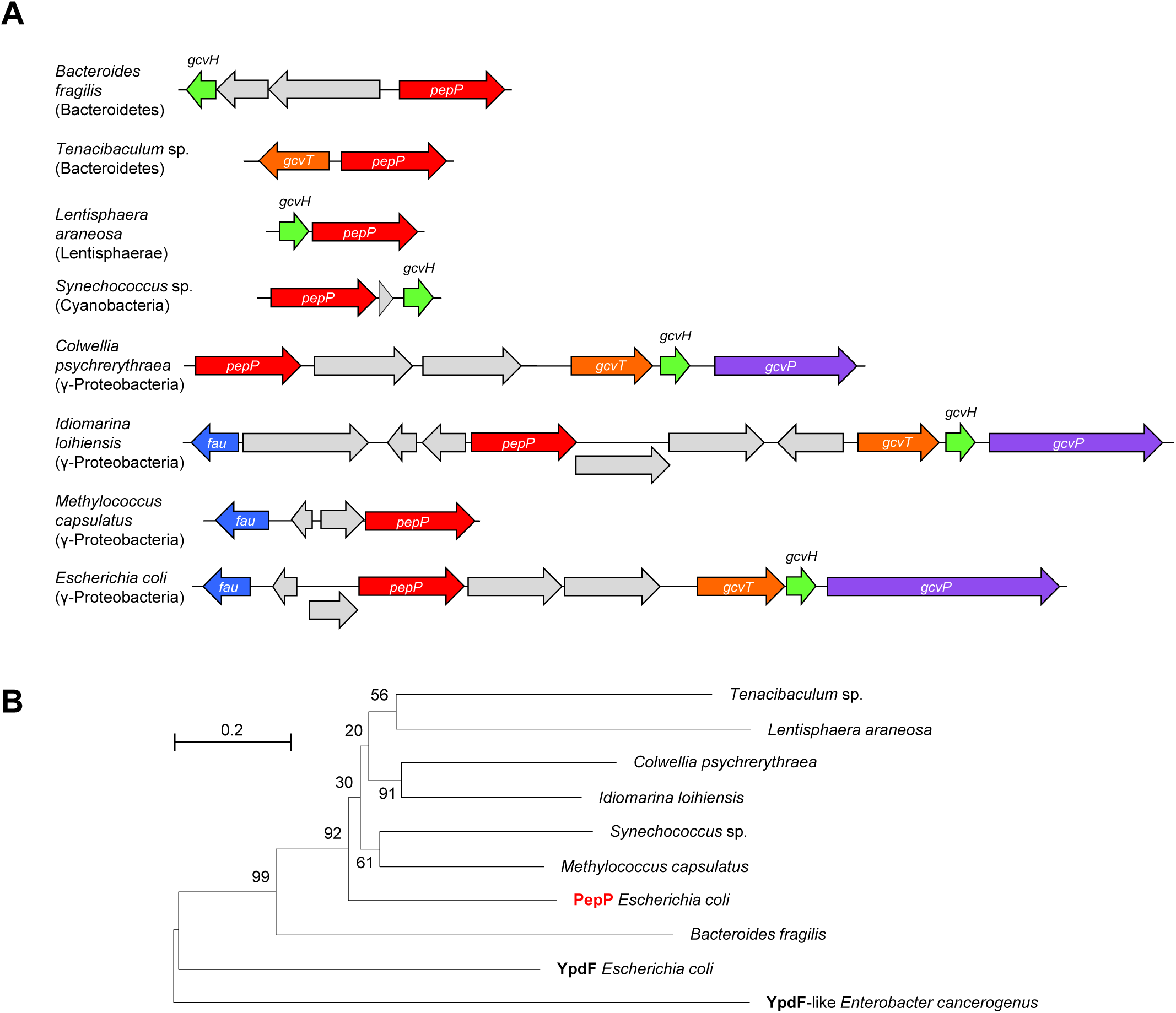
Bioinformatic analysis of PepP homologs from diverse bacteria. **(A)** PepP (Xaa-Pro aminopeptidase) homologs from four bacterial phyla cluster on the chromosome with genes encoding enzymes of one-carbon metabolism: 5-formyl-THF cyclo-ligase (*fau*) and the T, H, and P subunits of the glycine cleavage complex (*gcvT, gcvH*, and *gcvP*, respectively). Gray genes encode proteins whose function is either unrelated to one-carbon metabolism or unknown. **(B)** Neighbor-joining tree (500 bootstrap replicates) confirming that the proteins encoded by the *pepP* homologs above belong to the same clade as *E. coli* PepP, and not to other aminopeptidase clades (YpdF or YpdF-like clades). Bootstrap values are shown for branch support. Evolutionary distances are in units of amino acid substitutions per site.

### PepP contributes to formaldehyde resistance in *E. coli*

The above genomic evidence linking PepP with one-carbon metabolism led us to compare the HCHO resistance of an *E. coli pepP* deletion strain with that of the parent BW25113 strain (henceforth referred to as wild type). The Δ*pepP* strain was markedly more sensitive to HCHO at high concentrations (400– 500 μM) when cultured in liquid or solid medium (Figure 3A,B). To confirm that the *pepP* deletion causes the HCHO-sensitive phenotype (i.e., that no mutations elsewhere in the genome are involved) we complemented the Δ*pepP* strain with a plasmid-borne *E. coli pepP* gene (Fig. 3A). The methanol stabilizer present in the HCHO reagent (37% HCHO, 10-15% methanol) did not detectably affect the growth of the wild type or Δ*pepP* strains when added to plates at the concentration that would have accompanied the HCHO.

**Figure 3.**
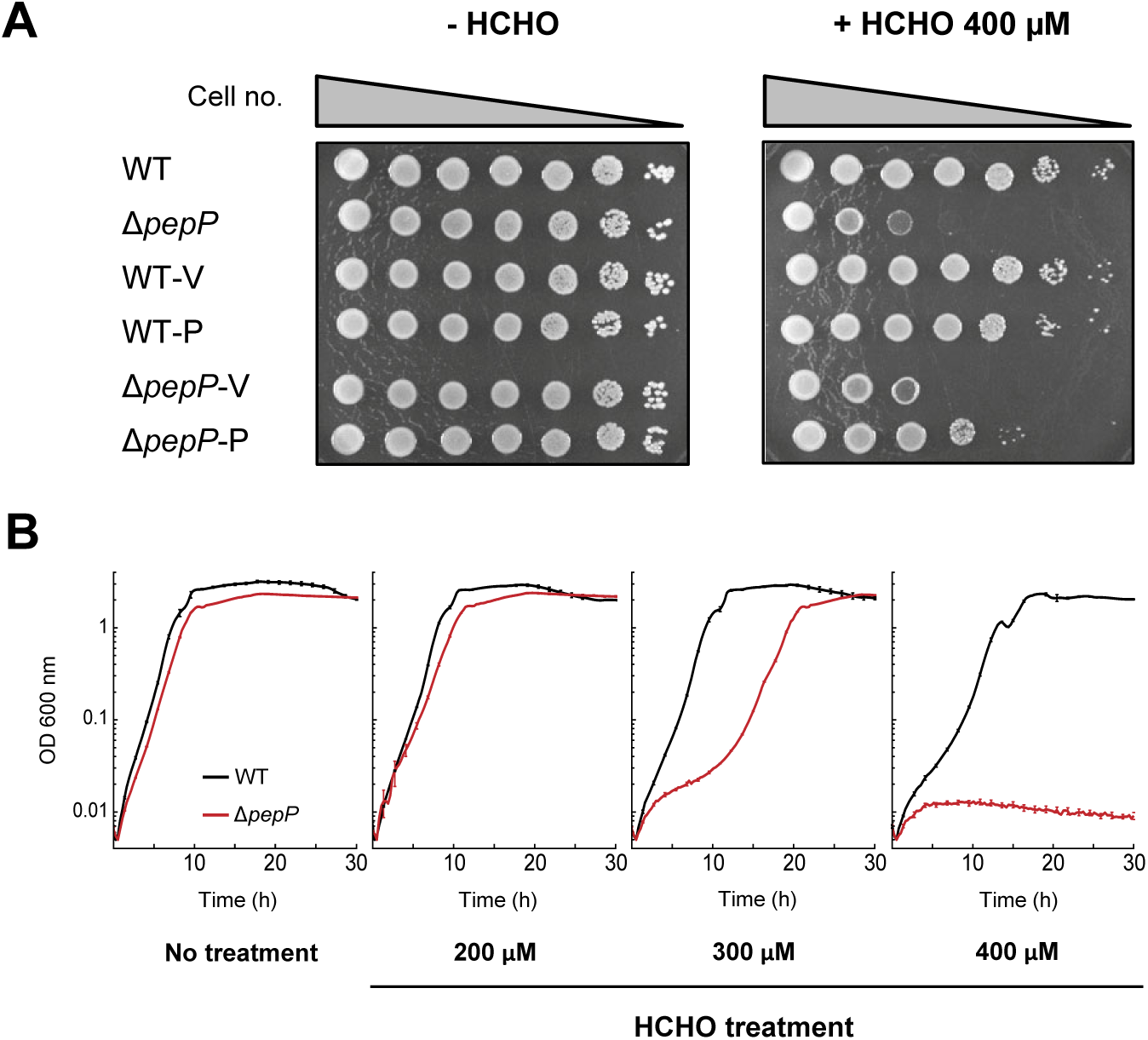
Growth assays implicate *E. coli* PepP in resistance to formaldehyde. **(A)** *E. coli* BW25113 wild type (WT) and Δ*pepP* strains harboring the pBAD24 vector alone (V) or containing *pepP* (P) were cultured on M9 minimal medium containing 0.4% glucose and 0.02% arabinose, minus or plus 400 μM HCHO. Overnight liquid cultures of each strain were ten-fold serially diluted and 3.5-μl aliquots were spotted on the plates. Images were captured after incubation at 37°C for 1. d. **(B)** Wild type and Δ*pepP* strains were cultured in 150 μl of M9 liquid medium containing 10 mM glucose and the indicated concentrations of HCHO. Data are means ± s.e. (*n* = 3).

To determine whether PepP impacts resistance to carbonyl compounds besides HCHO, we compared the growth of the parent and Δ*pepP* strains in the presence of inhibitory concentrations of acetaldehyde, glyoxal, or methylglyoxal. The Δ*pepP* deletant showed little or no increase in sensitivity to these carbonyls (Supplementary Figure S1), showing that PepP action is essentially specific to HCHO. We also tested whether the HCHO-sensitivity of the Δ*pepP* strain depends on the nature or concentration of the carbon source. Similar sensitivity was observed on eight diverse carbon sources, which included fermentable and non-fermentable substrates taken up by the phosphotransferase system or other carriers (Supplementary Figure S2). Sensitivity was also similar at various concentrations of glucose (Supplementary Figure S3). The HCHO-sensitivity of the Δ*pepP* strain is thus a robust phenotype.

### PepP does not deformylate *N*^6^-formyl lysine *in vitro* or *in vivo*

*E. coli* PepP (Xaa-Pro aminopeptidase, EC 3.4.11.9) cleaves the N-terminal amino acid from a wide range of peptides whose second residue is proline; the reaction catalyzed is thus X-Pro-Y_*n*_ → X + Pro-Y_*n*_ (where *n* can be up to ∼10) [30]. Because *N*^6^-formylation of lysine residues is a known HCHO-driven protein damage reaction [11,12] and because some peptidases have amidase activity [31,32], we tested whether PepP cleaves the amide bond in *N*^6^-formyl lysine (i.e. deformylates lysine) *in vitro* or *in vivo*. For *in vitro* tests, *N*^6^-formyl lysine and its N- and C-terminally blocked derivative Boc-*N*^6^-formyl Lys-AMC were incubated with recombinant *E. coli* PepP and a coupled spectrophotometric assay was used to measure formate release. No activity was detected with either substrate. The activity detection limit (0.6 nmol mg^−1^ protein min^−1^) was <0.01% of the activity of PepP against the canonical peptide substrate Ala-Pro-Ala (see below). To extend the study from model substrates to proteins *in vivo* we measured *N*^6^-formyl lysine levels in proteins extracted from wild type and Δ*pepP* cells before and after treatment with a nonlethal, bacteriostatic concentration of HCHO (Figure 4). HCHO-treated wild type and Δ*pepP* cells had modestly higher *N*^6^-formyl lysine levels than untreated cells, consistent with the known formylating activity of HCHO [11,12]. However, *N*^6^-formyl lysine levels in Δ*pepP* cells did not differ significantly from those in wild type cells either before HCHO treatment (when cells were growing) or after HCHO treatment (when cells were not growing). The lack of effect of deleting *pepP* agrees with the *in vitro* evidence that PepP does not have lysine-deformylating activity and hence confirms that this activity cannot explain the role of PepP in HCHO resistance.

**Figure 4.**
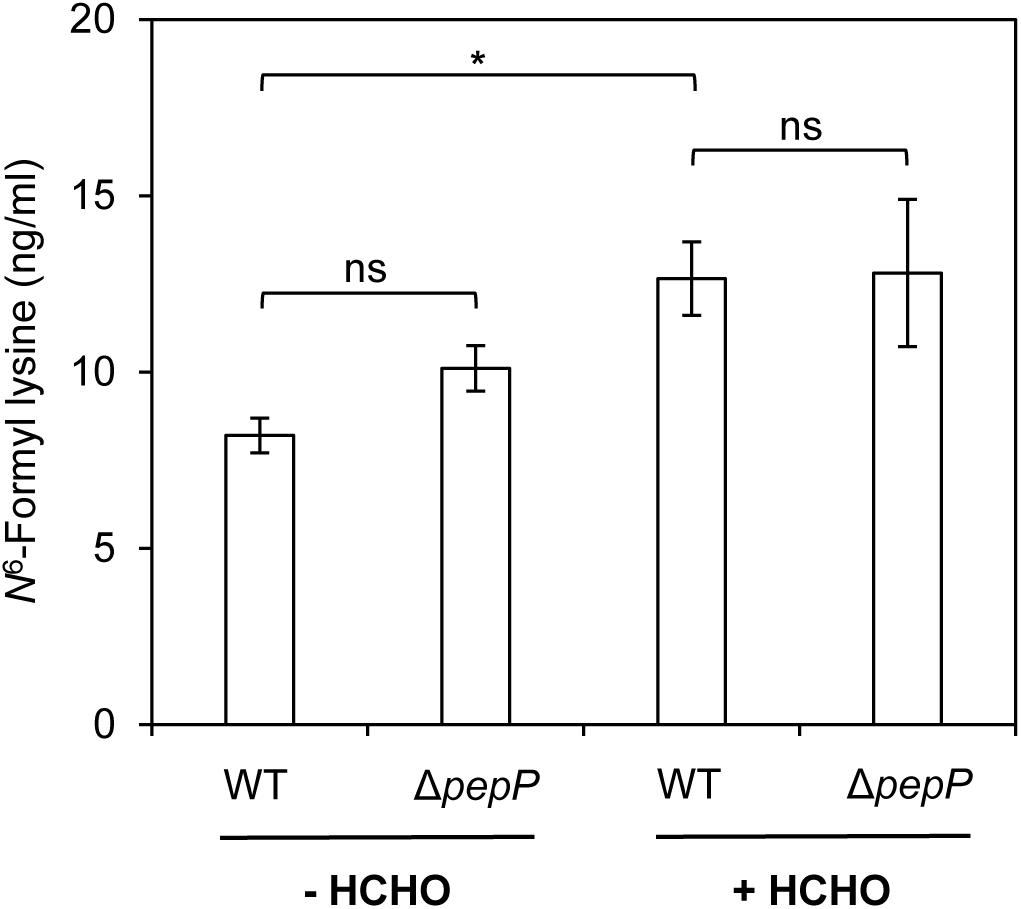
*N*^*6*^-Formyl lysine levels in proteins from wild type and Δ*pepP* strains. *E. coli* BW25113 wild type (WT) and Δ*pepP* strains were grown in M9 minimal medium containing 0.4% glucose until OD_600_ reached 0.4. Control (-HCHO) cells were then harvested; HCHO-treated (+ HCHO) cells were cultured for another 2 h after adding HCHO (final concentration 1 mM) and then harvested. Extracted proteins were hydrolyzed with *S. griseus* protease; the hydrolysate was analyzed by LC-MS for *N*^6^-formyl lysine. Data are means ± s.e. (*n* = 6). Significance was determined by Student’s *t*-test. \**P* <0.05; ns, non-significant.

### A deductive hypothesis connecting PepP to formaldehyde toxicity

It is not obvious how an aminopeptidase like PepP could impact HCHO toxicity unless it has substantial deformylase activity, which the above evidence indicates it does not. However, a plausible multi-part hypothesis can be proposed, based on five facts: (i) thioproline is a close analog of proline, with similar bond lengths and angles [33]; (ii) thioproline forms spontaneously *in vivo* from the condensation of HCHO and free or protein-bound cysteine [10,11]; (iii) thioproline is charged on tRNA^Pro^ and enters proteins [33,34]; (iv) PepP is specific for proline as the second residue in its peptide substrates [30]; and (v) free thioproline is detoxified via proline dehydrogenase (PutA)-mediated oxidation to 2,3-thiazoline-4-carboxylate followed by hydrolysis to cysteine plus formate [35,36]. Our hypothesis is as follows: HCHO buildup causes formation of thioproline-containing proteins whose degradation produces thioproline-containing peptides; these peptides are toxic; PepP is necessary for hydrolysis of these peptides, enabling release of free thioproline that is safely metabolized via PutA to cysteine and formate (Figure 5A). The core of the hypothesis (Figure 5A, box) is that thioproline peptides are toxic and that PepP is essential for their removal.

**Figure 5.**
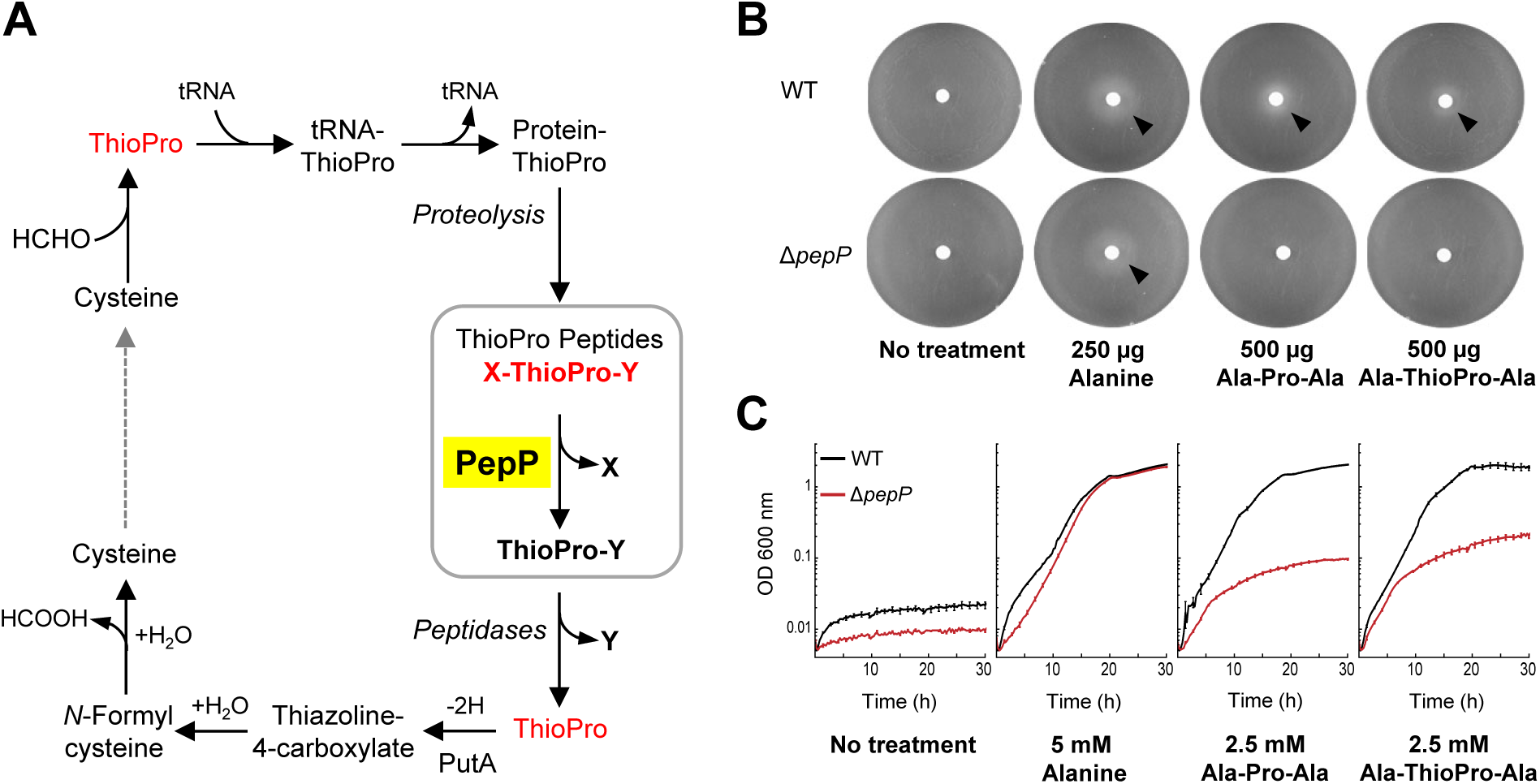
A hypothesis connecting PepP to formaldehyde toxicity and evidence supporting it. **(A)** Schematic representation of the hypothesis. Compounds inferred or known to be toxic are in red font. The core parts of the hypothesis – the toxicity of thioproline-containing peptides and their cleavage by PepP – are boxed in gray. ThioPro, thioproline; X and Y represent any amino acid residue. **(B)** Growth of lawns of BW25113 wild type (WT) and Δ*pepP* cells on plates of M9 minimal medium without N, plus 0.4% glucose. The central disc contained the additions indicated; alanine was included as a positive control. Arrowheads mark growth halos around the discs. **(A)** Growth of wild type and Δ*pepP* cells in liquid M9 minimal medium without N, plus 10 mM glucose and the additions indicated. Data are means ± s.e. (*n* = 3).

This hypothesis explains why deleting PepP does not increase sensitivity to acetaldehyde, glyoxal, or methylglyoxal (Supplementary Figure S1). Acetaldehyde reacts with free cysteine to form the thioproline analog 2-methylthioproline [37] but this amino acid is almost surely not charged to tRNA and incorporated into proteins [38], and acetaldehyde has not been reported to react with peptidyl-cysteine; hence there would be no 2-methylthioproline-containing peptides for PepP to cleave. Glyoxal and methylglyoxal cannot react with cysteine to form thiazolidine adducts (i.e. thioproline-type derivatives) [39], so again there would be no thioproline-like peptides for PepP to act on.

The hypothesis also makes two testable predictions, namely that: (i) PepP readily hydrolyzes peptides with the general formula X-thioproline-Y to X + thioproline-Y (X and Y being any amino acid); and (ii) deleting *pepP* increases sensitivity to supplied thioproline because this is incorporated into proteins [34] that are cleaved to peptides that are toxic. Our next step was to test these predictions.

### Experimental tests of hypothesis predictions

We first tested the activity of purified recombinant PepP against the thioproline-containing peptide Alathioproline-Ala using its natural counterpart Ala-Pro-Ala as a benchmark (Ala-Pro-Ala is known to be a good PepP substrate [30,40]). Activity was assayed via a coupled spectrophotometric assay that measured release of alanine. As pilot experiments showed similar activity against both peptides, we proceeded to a full kinetic characterization (Table 1 and Supplementary Figure S4). Our *K*_m_ and *k*_cat_ values for the natural substrate Ala-Pro-Ala were consistent with those previously reported for this peptide [40]. As predicted, Ala-thioproline-Ala was a good substrate, giving *K*_m_ and *k*_cat_ values that did not differ statistically from those for Ala-Pro-Ala. More generally, the *K*_m_ and *k*_cat_ values for Ala-thioproline-Ala were within the range reported for PepP acting on various peptides [30]. PepP thus acted on a model thioproline-containing peptide just as if it were a regular proline-containing peptide.

**Table 1.** Kinetic parameters of recombinant *E. coli* PepP. A coupled spectrophotometric assay was used. Values are means ± s.e. (*n* = 3). The *K*_m_ value for Ala-Pro-Ala is close to the reported value (0.77 mM) [40] and the *k*_cat_ value is consistent with that reported previously (85 s^-1^) [40], allowing for differences in assay pH and temperature. All values for Ala-thioproline-Ala fall in the 95% confidence intervals for Ala-Pro-Ala values, i.e. are statistically the same.

| Substrate | $V_{max}$ ($\mu\text{mol min}^{-1} \text{mg}^{-1}$) | $k_{cat}$ (s<sup>-1</sup>) | $K_m$ (mM) | $k_{cat}/K_m$ (s<sup>-1</sup> M<sup>-1</sup>) |
| --- | --- | --- | --- | --- |
| Ala-Pro-Ala | 16.4 ± 1.6 | 14.0 ± 1.3 | 1.10 ± 0.26 | 1.27 $\times 10^4$ |
| Ala-thioproline-Ala | 16.2 ± 0.9 | 13.8 ± 0.9 | 1.01 ± 0.15 | 1.37 $\times 10^4$ |

To confirm that PepP cleaves Ala-thioproline-Ala *in vivo* as well as *in vitro* we exploited the ability of *E. coli* to use alanine as sole nitrogen source [41]. We reasoned that if PepP is the sole or main peptidase that cleaves the N-terminal amino acid from X-thioproline-Y and X-proline-Y peptides, and if such peptides are not hydrolyzed by carboxypeptidase activity [42], then wild type *E. coli* should be able to use Ala-thioproline-Ala or Ala-Pro-Ala as sole nitrogen source and the Δ*pepP* strain should not. This proved to be the case (Figure 5B). Plates containing a lawn of cells and a central disc charged with either tripeptide gave growth halos around the disc with wild type cells but not with Δ*pepP* cells. Comparable results were obtained in liquid culture, i.e. the Δ*pepP* deletion strain lost most of its capacity to use Ala-thioproline-Ala or Ala-Pro-Ala as sole nitrogen source (Figure 5C). The residual growth of the Δ*pepP* deletant was presumably due to a low level of alanine release by another peptidase(s). Note that the similar growth patterns of wild type and Δ*pepP* cells on Ala-Pro-Ala or Ala-thioproline-Ala show that Ala-thioproline-Ala was not toxic at the concentration used (2.5 mM). Finally, we compared the responses of the wild type and Δ*pepP* strains to supplied thioproline. As predicted, the Δ*pepP* strain was more sensitive, becoming progressively more inhibited than wild type as the thioproline concentration was increased from 2 mM to 4 mM (Figure 6).

**Figure 6.**
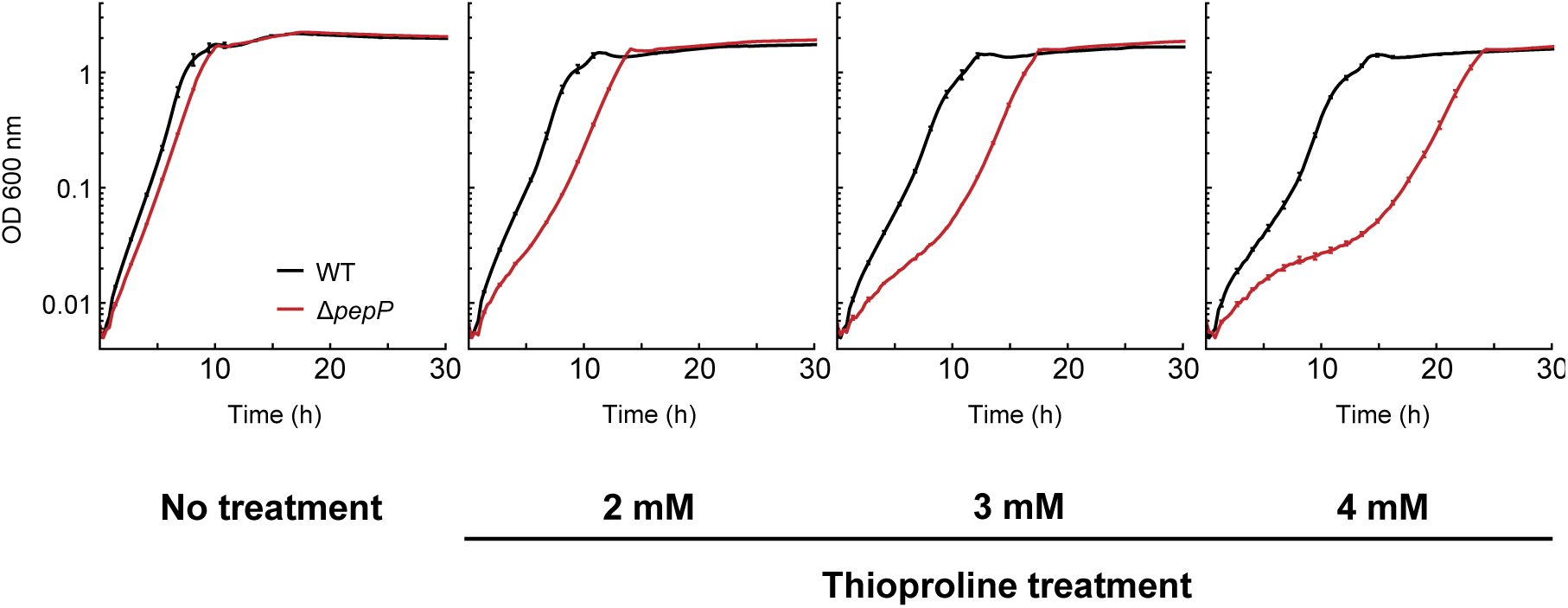
Deleting *pepP* increases sensitivity to supplied thioproline. BW25113 wild type (WT) and Δ*pepP* strains were cultured in 150 μl of M9 liquid medium containing 10 mM glucose and the indicated concentrations of thioproline. Data are means ± s.e. (*n* = 3).

### Ancillary experimental tests

Given the role of PutA in detoxifying thioproline (Figure 5A) [35], we compared the effect of deleting *putA* on sensitivity to HCHO to that of deleting *pepP*, using a concentration of HCHO (400 µM) that inhibits growth of the Δ*pepP* strain (Supplementary Figure S5). At this HCHO concentration, the Δ*putA* strain showed no growth defect. This result implies that the thioproline-containing peptides that PepP hydrolyzes are more toxic than thioproline itself. PepP thus appears to be more critical than PutA in the chain of reactions that cope with protein-bound thioproline (Figure 5A).

Because glutathione is ∼100-fold more abundant in *E. coli* than free cysteine [43,44] and reacts readily with HCHO to form *S*-(hydroxymethyl)glutathione and other adducts [45] we tested whether PepP can hydrolyze the adduct γ-Glu-thioproline-Gly. This compound could conceivably form by spontaneous reaction of HCHO with glutathione’s cysteine residue, although it is not one of the reported HCHO adducts of glutathione [45]. γ-Glu-thioproline-Gly was incubated with PepP and glutamate release was assayed by a coupled spectrophotometric procedure. No activity was detected; the detection limit (2 nmol mg^−1^ protein min^−1^) was <0.01% of the activity of PepP against Ala-Pro-Ala.

### The Keio collection Δ*yhbO* strain carries a *pepP* nonsense mutation

In the course of this work we found by genome sequencing that the Keio collection Δ*yhbO* strain carries a nonsense mutation in the *pepP* ORF that changes codon 285 from CAG (glutamine) to TAG (stop). The resulting truncated PepP protein is evidently non-functional because the Keio Δ*yhbO* strain had the same HCHO sensitivity as the Δ*pepP* strain (Supplementary Figure S6). A Δ*yhbO* strain lacking the *pepP* nonsense mutation (constructed by P1 phage transduction from the BW25113 Keio parent strain) showed no increase in HCHO sensitivity (Supplementary Figure S6), confirming that the HCHO sensitivity of the *pepP-*Δ*yhbO* strain is due solely to the PepP defect.

## Conclusions

The core parts of our hypothesis connecting the Xaa-Pro aminopeptidase PepP with HCHO resistance are (i) that thioproline-containing peptides (at least those with thioproline as second residue) are toxic, and (ii) that PepP can cleave Xaa-thioproline peptides (Figure 5A). Part (ii) is validated by the evidence presented above. Part (i) is a strong inference – and will likely remain so because it is so challenging to test. With thioproline at position two, there are 20^2^ (400) possible tripeptides, 20^3^ (8,000) possible tetrapeptides, and so forth up to 20^9^ (512 billion) possible decapeptides. Assuming, as for other peptides [46,47], that toxicity is restricted to a tiny fraction of the total sequence space, it would be necessary to screen an intractable number of peptides of various sizes and sequences in order to identify the few that are toxic. Moreover, these peptides would have to be supplied exogenously but might not be taken up because bacterial peptide transporters have various degrees of substrate specificity [48], or might be hydrolyzed before uptake by peptidases released by dead cells [49].

Although toxicity testing of thioproline-containing peptides is infeasible, it is interesting to ask what the basis of their toxicity could be. We suggest the following general explanation, based on the findings that certain non-natural oligopeptides inhibit *E. coli* growth [50] and that thioproline-containing polypeptides can differ greatly in physicochemical properties from their natural proline-containing counterparts [34]. Perhaps certain thioproline-containing peptides simply differ enough from their harmless natural analogs to be toxic via the same undefined mechanisms as other alien oligopeptides [50].

In sum, our genetic and biochemical findings on PepP provide strong evidence that the formation of thioproline-containing peptides is a substantial driver of HCHO toxicity. From an engineering standpoint, this evidence makes PepP a potential damage-control component [17] for use in designing and building pathways in which HCHO is an intermediate (Figure 1). Although the damage-control pathway involving PepP (Figure 5A) ultimately converts HCHO to formate, the flux via this pathway is constrained by the rate of proteolysis and is hence almost certainly too low to constitute a major drain on the HCHO pool. From a physiological standpoint, the evidence that PepP deals with a toxic consequence of HCHO formation – which happens in normal metabolism [1-3] – raises a question: Is cleaving thioproline peptides the primary role of PepP? If so, the well-characterized activity of PepP against proline peptides would be of secondary importance. That the Δ*pepP* strain has no growth defect on minimal medium (Figure 3A,B) fits with this possibility. So too does the inference [51] that *Lactococcus lactis* PepP does not have a major role in nitrogen nutrition and probably has some other function.

## Acknowledgments

We thank Laszlo N. Csonka for his insightful advice on bacterial genetics, and Michael J. Ziemak for his contribution to advancing this work.

## Author Contribution

A.D.H., A.B.-E., J.A.P., H.H., and M.A.W. conceived and designed the research; J.A.P. and H.H. performed experiments with supervision from A.D.H. and A.B.-E.; Q.L. and S.D.B. synthesized substrates; J.S.F. and O.F. carried out metabolomic analyses; A.D.H. made computational gene clustering analyses; J.A.P. and A.D.H. wrote the article.

## Funding

This research was supported by U.S. National Science Foundation awards MCB-1611711 to A.D.H. and MCB-1611846 to O.F., by Hatch Project FLA-HOS-005796, and by an endowment from the C.V. Griffin Sr. Foundation. A.B.-E. and H.H. are funded by the Max Planck Society. H.H. is also funded by the China Scholarship Council.

## Competing Interests

The Authors declare that there are no competing interests associated with the manuscript. A.B.-E. is co-founder of b.fab, which aims to commercialize microbial growth on C_1_ compounds. b.fab was not involved in this study and did not fund it.

## Abbreviations

HCHO: formaldehyde
THF: tetrahydrofolate

## References

1. Chen, N.H., Djoko, K.Y., Veyrier, F.J. and McEwan, A.G. (2016) Formaldehyde stress responses in bacterial pathogens. Front. Microbiol. 7, 257

2. Hanson, A.D. and Roje, S. (2001) One-carbon metabolism in higher plants. Annu. Rev. Plant Physiol. Plant Mol. Biol. 52, 119–137

3. Dringen, R., Brandmann, M., Hohnholt, M.C. and Blumrich, E.M. (2015) Glutathione-dependent detoxification processes in astrocytes. Neurochem Res. 40, 2570–2582

4. Cotton, C.A., Claassens, N.J., Benito-Vaquerizo, S. and Bar-Even, A. (2019) Renewable methanol and formate as microbial feedstocks. Curr. Opin. Biotechnol. 62, 68–180

5. Yishai, O., Lindner, S.N., Gonzalez de la Cruz, J., Tenenboim, H. and Bar-Even A. (2016) The formate bio-economy. Curr. Opin. Chem. Biol. 35, 1–9

6. Kim SJ, Yoon J, Im DK, Kim YH, Oh MK (2019) Adaptively evolved *Escherichia coli* for improved ability of formate utilization as a carbon source in sugar-free conditions. Biotechnol. Biofuels 12, 207

7. Schroer, K., Luef, K.P., Hartner, F.S., Glieder, A. and Pscheidt, B. (2010) Engineering the *Pichia pastoris* methanol oxidation pathway for improved NADH regeneration during whole-cell biotransformation. Metab. Eng. 12, 8–17

8. Metz, B., Kersten, G.F., Hoogerhout, P., Brugghe, H.F., Timmermans, H.A., de Jong, A. et al. (2004) Identification of formaldehyde-induced modifications in proteins: reactions with model peptides. J. Biol. Chem. 279, 6235–6243

9. Hoffman, E.A., Frey, B.L., Smith, L.M. and Auble, D.T. (2015) Formaldehyde crosslinking: a tool for the study of chromatin complexes. J. Biol. Chem. 290, 26404–26411

10. Kamps, J.J.A.G., Hopkinson, R.J., Schofield, C.J. and Claridge, T.D.W. (2019) How formaldehyde reacts with amino acids. Commun. Chem. 2, 126

11. Liu, J., Chan, K.K. and Chan, W. (2016) Identification of protein thiazolidination as a novel molecular signature for oxidative stress and formaldehyde exposure. Chem. Res. Toxicol. 29, 1865–1871

12. Wisniewski, J.R., Zougman, A. and Mann, M. (2008) *N*6-Formylation of lysine is a widespread post-translational modification of nuclear proteins occurring at residues involved in regulation of chromatin function. Nucleic Acids Res. 36, 570–577

13. Van Spanning, R.J.M., de Vries, S. and Harms, N. (2000) Coping with formaldehyde during C1 metabolism of *Paracoccus denitrificans*. J. Mol. Catal. B: Enzym. 8, 37–50

14. Jakobsen, Ø.M., Benichou, A., Flickinger, M.C., Valla, S., Ellingsen, T.E. and Brautaset, T. (2006) Upregulated transcription of plasmid and chromosomal ribulose monophosphate pathway genes is critical for methanol assimilation rate and methanol tolerance in the methylotrophic bacterium *Bacillus methanolicus*. J. Bacteriol. 188, 3063–3072

15. Seidel, J., Klockenbusch, C. and Schwarzer, D. (2016) Investigating deformylase and deacylase activity of mammalian and bacterial sirtuins. ChemBioChem 17, 398–402

16. Bouzon, M., Perret, A., Loreau, O., Delmas, V., Perchat, N., Weissenbach, J. et al. (2017) A synthetic alternative to canonical one-carbon metabolism. ACS Synth. Biol. 6, 1520–1533

17. Sun, J., Jeffryes, J.G., Henry, C.S., Bruner, S.D. and Hanson, A.D. (2017) Metabolite damage and repair in metabolic engineering design. Metab. Eng. 44, 150–159

18. Szklarczyk, D., Gable, A.L., Lyon, D., Junge, A., Wyder, S., Huerta-Cepas, J. et al. (2019) STRING v11: protein-protein association networks with increased coverage, supporting functional discovery in genome-wide experimental datasets. Nucleic Acids Res. 47, D607–D613

19. Overbeek, R., Begley, T., Butler, R.M., Choudhuri, J. V., Chuang, H.Y., Cohoon, M. et al. (2005) The subsystems approach to genome annotation and its use in the project to annotate 1000 genomes. Nucleic Acids Res. 33, 5691–5702

20. Tamura, K., Stecher, G., Peterson, D., Filipski, A. and Kumar, S. (2013) MEGA6: Molecular Evolutionary Genetics Analysis version 6.0. Mol. Biol. Evol. 30, 2725–2729

21. McClure, J.J., Inks, E.S., Zhang, C., Peterson, Y.K., Li, J., Chundru, K. et al. (2017) Comparison of the deacylase and deacetylase activity of zinc-dependent HDACs. ACS Chem. Biol. 12, 1644–1655

22. Baba, T., Ara, T., Hasegawa, M., Takai, Y., Okumura, Y., Baba, M. et al. (2006) Construction of *Escherichia coli* K-12 in-frame, single-gene knockout mutants: the Keio collection. Mol. Syst. Biol. 2, 2006.0008

23. Thomason, L.C., Costantino, N. and Court, D.L. (2007) *E. coli* genome manipulation by P1 transduction. Curr. Protoc. Mol. Biol. 79, 1.17.1-1.17.8

24. Quan, J. and Tian, J. (2009) Circular polymerase extension cloning of complex gene libraries and pathways. PLoS ONE 4, e6441

25. Bradford, M.M. (1976) A rapid and sensitive method for the quantitation of microgram quantities of protein utilizing the principle of protein-dye binding. Anal. Biochem. 72, 248–254

26. de Crécy-Lagard, V., Haas, D. and Hanson, A.D. (2018) Newly-discovered enzymes that function in metabolite damage-control. Curr. Opin. Chem. Biol. 47, 101–108

27. Kallen, R.G. and Jencks, W.P. (1966) The mechanism of the condensation of formaldehyde with tetrahydrofolic acid. J. Biol. Chem. 241, 5851–5863

28. Douce, R., Bourguignon, J., Neuburger, M. and Rébeillé, F. (2001) The glycine decarboxylase system: a fascinating complex. Trends Plant Sci. 6, 167–176

29. Stover, P. and Schirch, V. (1990) Serine hydroxymethyltransferase catalyzes the hydrolysis of 5,10-methenyltetrahydrofolate to 5-formyltetrahydrofolate. J. Biol. Chem. 265, 14227–14233

30. Yoshimoto, T., Orawski, A.T. and Simmons, W.H. (1994) Substrate specificity of aminopeptidase P from *Escherichia coli*: comparison with membrane-bound forms from rat and bovine lung. Arch. Biochem. Biophys. 311, 28–34

31. Bompard-Gilles, C., Villeret, V., Davies, G.J., Fanuel, L., Joris, B., Frère, J.M. et al. (2000) A new variant of the Ntn hydrolase fold revealed by the crystal structure of L-aminopeptidase D-ala-esterase/amidase from *Ochrobactrum anthropi*. Structure 8, 153–162

32. Komeda, H., Hariyama, N. and Asano, Y. (2006) L-Stereoselective amino acid amidase with broad substrate specificity from *Brevundimonas diminuta*: characterization of a new member of the leucine aminopeptidase family. Appl. Microbiol. Biotechnol. 70, 412–421

33. Budisa, N., Minks, C., Medrano, F.J., Lutz, J., Huber, R. and Moroder, L. (1998) Residue-specific bioincorporation of non-natural, biologically active amino acids into proteins as possible drug carriers: structure and stability of the *per*-thiaproline mutant of annexin V. Proc. Natl. Acad. Sci. USA 95, 455–459

34. Liu, J., Hao, C., Wu, L., Madej, D., Chan, W. and Lam, H. (2020) Proteomic analysis of thioproline misincorporation in *Escherichia coli*. J. Proteomics 210, 103541

35. Unger, L. and DeMoss, R.D. (1966) Metabolism of a proline analogue, L-thiazolidine-4-carboxylic acid, by *Escherichia coli*. J. Bacteriol. 91, 1564–1569

36. Jeelani, G., Sato, D., Soga, T., Watanabe, H. and Nozaki, T. (2014) Mass spectrometric analysis of L-cysteine metabolism: physiological role and fate of L-cysteine in the enteric protozoan parasite *Entamoeba histolytica*. mBio 5, e01995

37. Liu, J., Meng, X. and Chan, W. (2016) Quantitation of thioprolines in grape wine by isotope dilution-liquid chromatography-tandem mass spectrometry. J. Agric. Food Chem. 64, 1361–1366

38. Saravanan Prabhu, N., Ayyadurai, N., Deepankumar, K., Chung, T. Lim, D. and Yun, H. (2012) Reassignment of sense codons: Designing and docking of proline analogs for *Escherichia coli* prolyl-tRNA synthetase to expand the genetic code. J. Mol. Catal. B 78, 57–64

39. Jeffryes, J.G., Colastani, R.L., Elbadawi-Sidhu, M., Kind, T., Niehaus, T.D., Broadbelt, L.J. et al. (2015) MINEs: open access databases of computationally predicted enzyme promiscuity products for untargeted metabolomics. J. Cheminform. 7, 44

40. Graham, S.C., Lilley, P.E., Lee, M., Schaeffer, P.M., Kralicek, A.V., Dixon, N.E. et al. (2006) Kinetic and crystallographic analysis of mutant *Escherichia coli* aminopeptidase P: insights into substrate recognition and the mechanism of catalysis. Biochemistry 45, 964–975

41. Varricchio, F. (1969) Control of glutamate dehydrogenase synthesis in *Escherichia coli*. Biochim. Biophys. Acta 177, 560–564

42. Yaron, A., Mlynar, D. and Berger, A. (1972) A dipeptidocarboxypeptidase from *E. coli*. Biochem. Biophys. Res. Commun. 47, 897–902

43. Wheldrake, J.F. (1967) Intracellular concentration of cysteine in *Escherichia coli* and its relation to repression of the sulphate-activating enzymes. Biochem. J. 105, 697–699

44. Smirnova, G.V., Tyulenev, A.V., Bezmaternykh, K.V., Muzyka, N.G., Ushakov, V.Y. and Oktyabrsky, O.N. (2019) Cysteine homeostasis under inhibition of protein synthesis in *Escherichia coli* cells. Amino Acids 51, 1577–1592

45. Hopkinson, R.J., Barlow, P.S., Schofield, C.J. and Claridge, T.D.W. (2010) Studies on the reaction of glutathione and formaldehyde using NMR. Org. Biomol. Chem. 8, 4915–4920

46. Gupta, S., Kapoor, P., Chaudhary, K., Gautam, A., Kumar, R., Open Source Drug Discovery Consortium et al. (2013) *In silico* approach for predicting toxicity of peptides and proteins. PLoS One 8, e73957

47. Wang, G., Li, X. and Wang, Z. (2015) APD3: the antimicrobial peptide database as a tool for research and education. Nucleic Acids Res. 44, D1087–D1093

48. Garai, P., Chandra, K. and Chakravortty, D. (2017) Bacterial peptide transporters: Messengers of nutrition to virulence. Virulence 8, 297–309

49. Lazdunski, A.M. (1989) Peptidases and proteases of *Escherichia coli* and *Salmonella typhimurium*. FEMS Microbiol. Rev. 5, 265–276

50. Walker, J.R., Roth, J.R. and Altman, E. (2001) An *in vivo* study of novel bioactive peptides that inhibit the growth of *Escherichia coli*. J. Pept. Res. 58, 380–388

51. Matos, J., Nardi, M., Kumura, H. and Monnet, V. (1998) Genetic characterization of *pepP*, which encodes an aminopeptidase P whose deficiency does not affect *Lactococcus lactis* growth in milk, unlike deficiency of the X-prolyl dipeptidyl aminopeptidase. Appl. Environ. Microbiol. 64, 4591–4595

