## Supplemental materials for "Thioproline formation as a driver of formaldehyde toxicity in *Escherichia coli*"

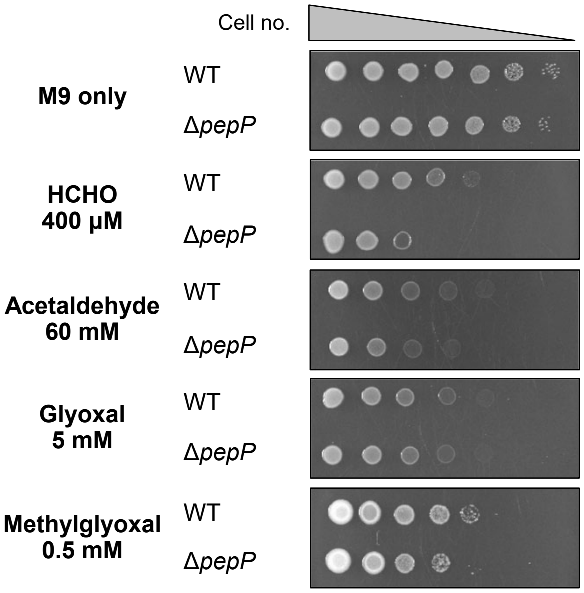


**Figure S1. Deletion of *pepP* has little or no effect on sensitivity to other small carbonyls.**

*E. coli* BW25113 (WT) and its Δ*pepP* derivative were cultured on M9 minimal medium containing 0.4% glucose and the indicated concentrations of HCHO, acetaldehyde, glyoxal, or methylglyoxal. Overnight liquid cultures of each strain were ten-fold serially diluted and 3.5-µl aliquots were spotted on the plates. Images were captured after incubation at 37°C for 1.5 d.

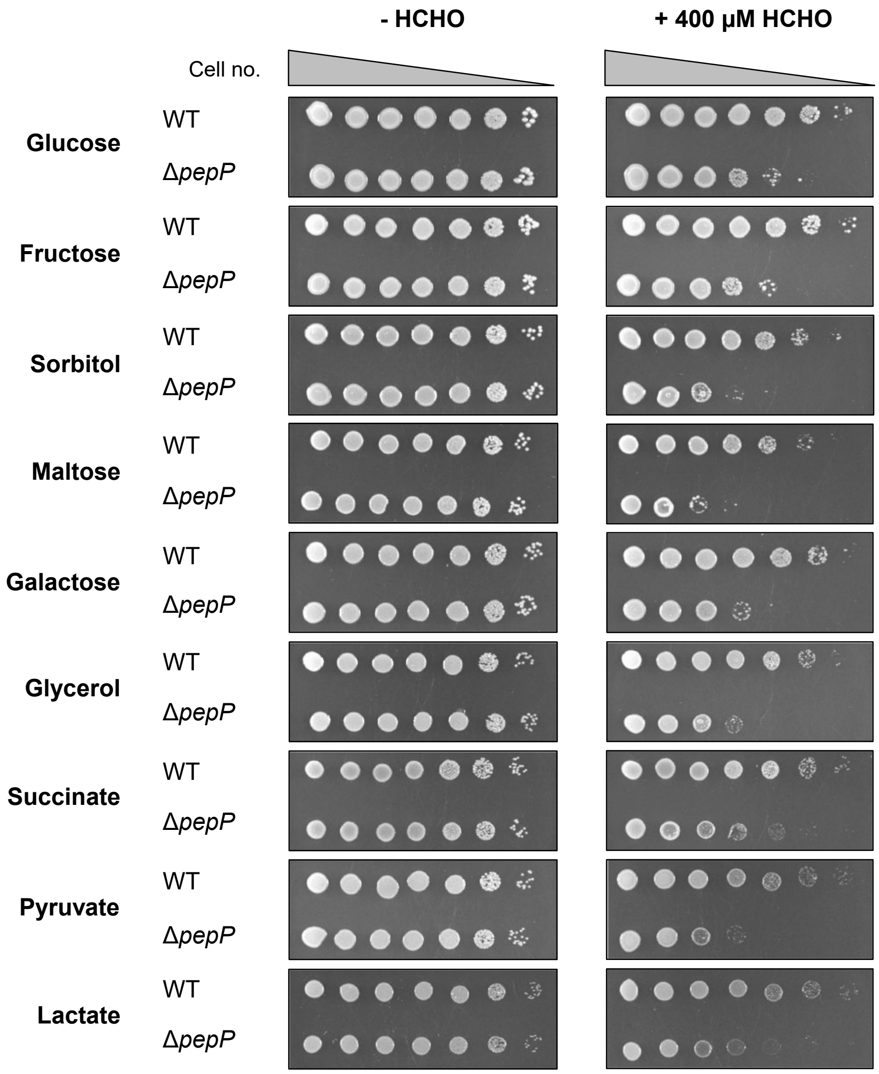


**Figure S2. HCHO-sensitivity of the Δ*pepP* strain is independent of carbon source.**

*E. coli* BW25113 (WT) and its Δ*pepP* derivative were cultured on M9 minimal medium containing 0.4% of the nine carbon sources shown, minus or plus 400 µM HCHO. Overnight liquid cultures of each strain were ten-fold serially diluted and 3.5-µl aliquots were spotted on the plates. Images were captured after incubation at 37°C for 1.5 d or, for plates containing lactate or pyruvate, 2 d.

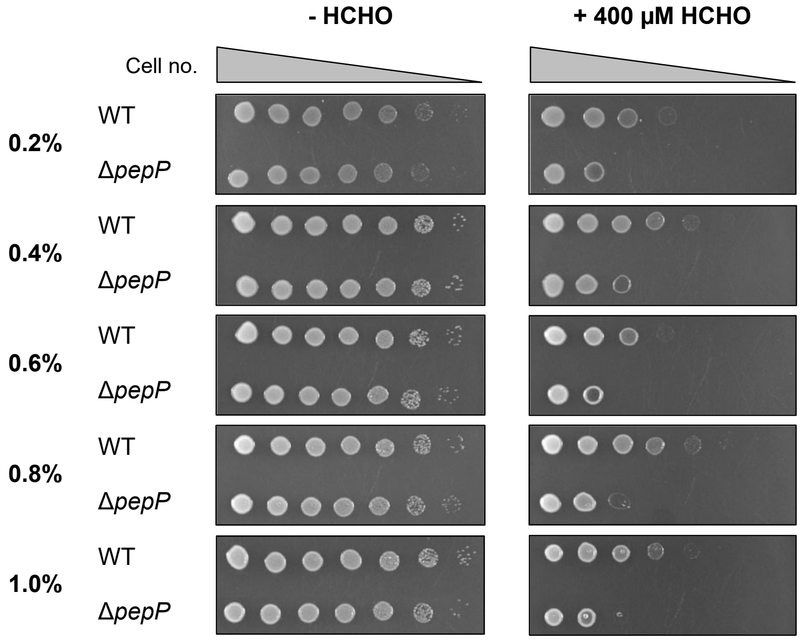


**Figure S3. HCHO-sensitivity of the Δ*pepP* strain is independent of glucose concentration.**

*E. coli* BW25113 (WT) and its Δ*pepP* derivative were cultured on M9 minimal medium containing glucose at the indicated concentrations, minus or plus 400 µM HCHO. Overnight liquid cultures of each strain were ten-fold serially diluted and 3.5-µl aliquots were spotted on the plates. Images were captured after incubation at 37°C for 1.5 d.

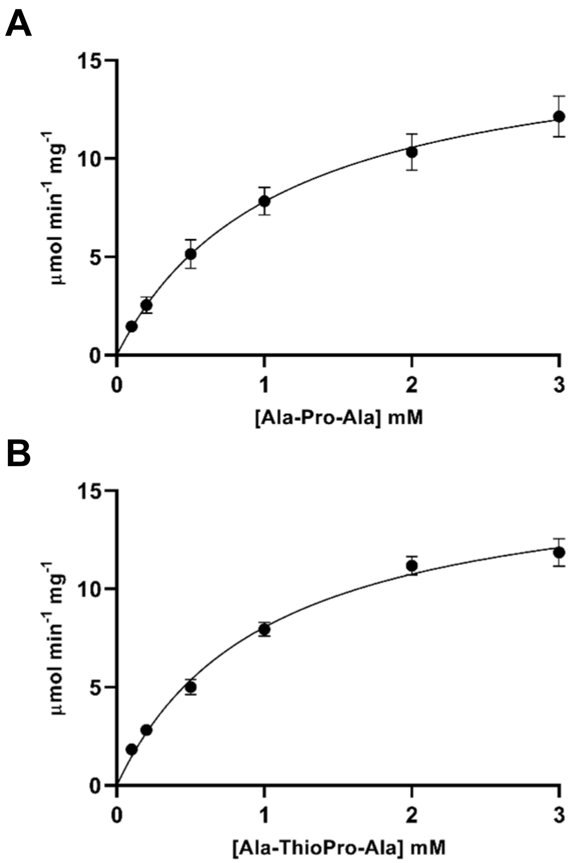


**Figure S4. Primary data from kinetic experiments.**

$K_m$ and $V_{max}$ values were determined by varying the concentration of Ala-Pro-Ala (**A**) or Ala-thioproline-Ala (**B**) in coupled spectrophotometric assays containing 50 mM Tris-HCl, pH 8.8, 0.5 mM $MnCl_2$, 1 mM NAD, 0.05 units of alanine dehydrogenase, and 2 μg PepP. The data were analyzed by nonlinear regression using GraphPad Prism. Data are means ± s.e. (*n* = 3).

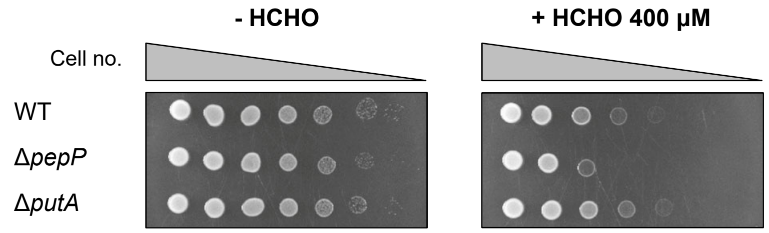


**Figure S5. Growth assays indicate that *E. coli* PutA does not contribute to HCHO resistance.**

*E. coli* BW25113 (WT), Δ*pepP* and Δ*putA* strains were cultured on M9 minimal medium containing 0.4% glucose, minus or plus 400 μM HCHO. Overnight liquid cultures of each strain were ten-fold serially diluted and 3.5-μl aliquots were spotted on the plates. Images were captured after incubation at 37°C for 1.5 d.

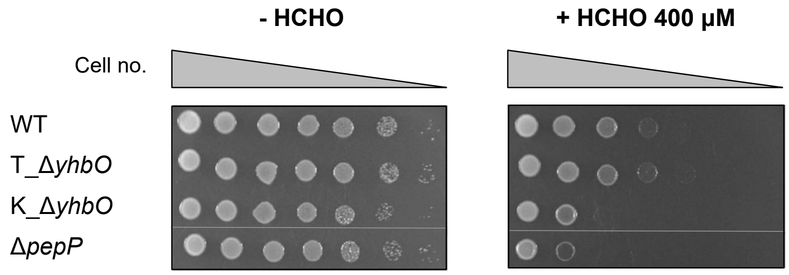


**Figure S6. A *pepP* mutation explains the formaldehyde sensitivity of the Keio Δ*yhbO* strain.**

*E. coli* BW25113 (WT), Keio Δ*yhbO* (K_Δ*yhbO*), P1 phage-transduced Δ*yhbO* (T_Δ*yhbO*), and Keio Δ*pepP* strains were cultured on M9 minimal medium containing 0.4% glucose, minus or plus 400 µM HCHO. Overnight liquid cultures of each strain were ten-fold serially diluted and 3.5-µl aliquots were spotted on the plates. Images were captured after incubation at 37°C for 1.5 d.

**Table S1. Primers for PCR amplification.**

| Primer | Name | Sequence |
|---|---|---|
| | | ***E. coli* Δ*pepP* complementation** |
| 1 | FP_pepP_EcoRI | AAAGAATTCACCATGAGTGAGATATCCCGGC |
| 2 | RP_pepP_SalI | TTTGTCGACTCATTGCTTTCTCGCAGC |
| | | **PepP protein expression** |
| 3 | FP_pepP_NC-term | ATGAGTGAGATATCCCG |
| 4 | RP_pepP_N-term | CTTTGTTAGCAGCCGGATCATTGCTTTCTCGC |
| 5 | RP_p15_N-term | CGGGATATCTCACTCATGCCGCTGCTGTGATGATGATG |
| 6 | FP_p15_NC-term | TCCGGCTGCTAACAAAGC |

**Table S2. MRM parameters for each targeted metabolite.**

Q1 and Q3 refer to the mass transitions monitored for each analyte. RT is the chromatographic retention time in minutes, DP, EP, CE, and CXP stand for declustering potential, entrance potential, collision energy, and cell exit potential respectively, and are measured in volts.

| Compound | Q1 (Da) | Q3 (Da) | RT (min) | DP | EP | CE | CXP |
|---|---|---|---|---|---|---|---|
| Lysine | 147.1 | 130.1 | 9.67 | 6 | 10 | 21 | 12 |
| $N^6$-formyl-formyl lysine | 175.1 | 112.1 | 8.36 | 26 | 10 | 19 | 12 |
| Cystine | 241 | 152 | 10.04 | 31 | 10 | 17 | 16 |
| D5 Threonine (IS) | 125.1 | 79.1 | 8.44 | 16 | 10 | 15 | 10 |
| D8 Lysine (IS) | 155.2 | 138.2 | 9.67 | 31 | 10 | 13 | 14 |
